# Direct, quantitative, and comprehensive analysis of tRNA acylation using intact tRNA liquid-chromatography mass-spectrometry

**DOI:** 10.1101/2023.07.14.549096

**Authors:** Riley Fricke, Isaac Knudson, Alanna Schepartz

## Abstract

Aminoacyl-tRNA synthetases (aaRSs) provide the functional and essential link between the sequence of an mRNA and the protein it encodes. aaRS enzymes catalyze a two-step chemical reaction that acylates specific tRNAs with a cognate α-amino acid. In addition to their role in translation, acylated tRNAs contribute to non-ribosomal natural product biosynthesis and are implicated in multiple human diseases. From the standpoint of synthetic biology, the acylation of tRNAs with a non-canonical α-amino acid (ncAA) or more recently, a non-α-amino acid monomer (nαAA) is a critical first step in the incorporation of these monomers into proteins, where they can be used for fundamental and applied science. These endeavors all demand an understanding of aaRS activity and specificity. Although a number of methods to monitor aaRS function *in vitro* or *in vivo* have been developed, many evaluate only the first step of the two-step reaction, require the use of radioactivity, or are slow, difficult to generalize, or both. Here we describe an LC-MS assay that rapidly, quantitatively, and directly monitors aaRS activity by detecting the intact acyl-tRNA product. After a simple tRNA acylation reaction workup, acyl- and non-acyl-tRNA molecules are resolved using ion-pairing reverse phase chromatography and their exact masses are determined using high-resolution time-of-flight mass spectrometry. The intact tRNA assay we describe is fast, simple, and quantifies reaction yields as low as 0.23%. The assay can also be employed on tRNAs acylated with flexizyme to detect products that are undetectable using standard techniques. The protocol requires basic expertise in molecular biology, mass spectrometry, and RNAse-free techniques.

## Introduction

Aminoacyl-tRNA synthetases provide the functional and essential link between the sequence of an mRNA and the protein it encodes. By physically linking an α-amino acid to a tRNA carrying the appropriate anticodon, aaRS enzymes both preserve and translate the genetic code. The chemical reaction catalyzed by an aaRS enzyme proceeds through two distinct steps. In the first step, an α-amino acid is activated for acyl exchange upon conversion into an aminoacyl-adenylate (aminoacyl-AMP). This reaction requires a single equivalent of ATP and releases inorganic pyrophosphate as a byproduct. In the second step, the activated aminoacyl-AMP acylates the 2’ or 3’ hydroxyl of the 3’ terminal adenosine of a tRNA co-substrate. This second step produces the final aminoacyl-tRNA product and releases AMP. (Fig. 1)^1^.

**Figure 1.**
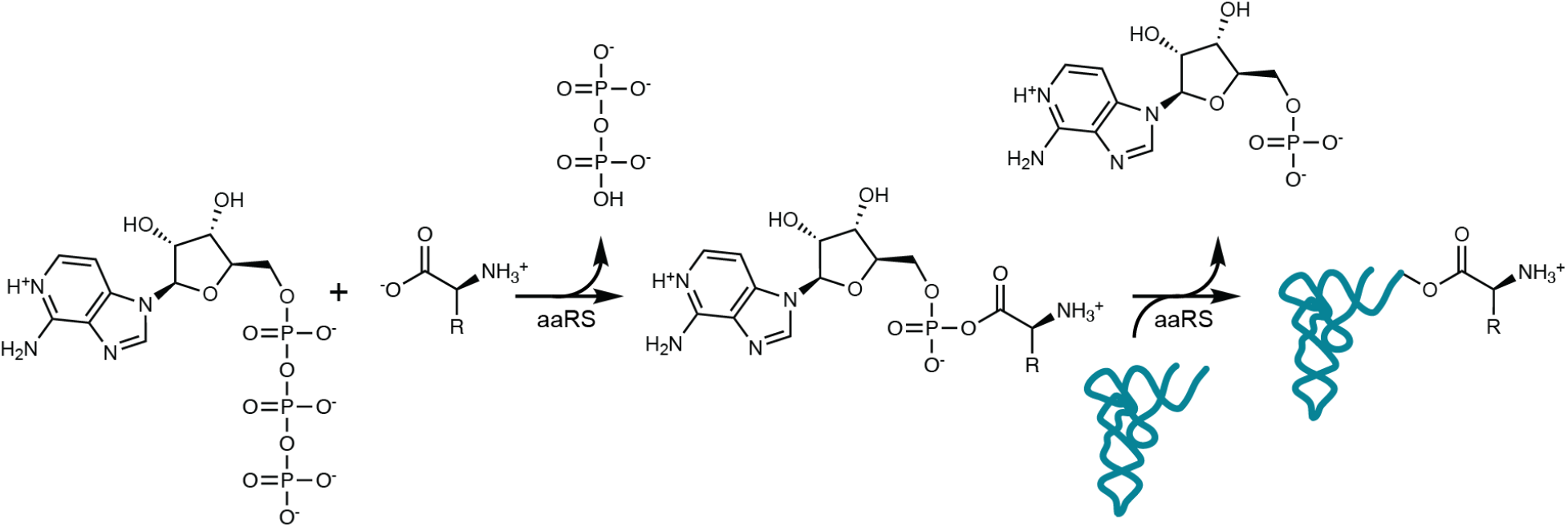
Two-step reaction catalyzed by aaRS enzymes. Some aaRS enzymes catalyze acylation of the 2’-hydroxyl group as opposed to the 3’-hydroxyl group as shown; isomerization between 2’ and 3’-isomers occurs rapidly^2^.

Aminoacyl-tRNA synthetases have been studied for decades because of their fundamental role in protein synthesis^1,3^. More recently they have gained additional attention because of their role in a variety of human diseases including neurodegenerative disorders, mitochondrial myopathies, HIV, Parkinson’s disease, autoimmune disorders, cancer, and diabetes.^3,4^ Aminoacyl-tRNA synthetases, their tRNA substrates, and acyl-tRNA products also play critical roles in natural product biosynthetic pathways^5^. Over the last three decades, aaRS enzymes have been studied widely and engineered to support the technology known as genetic code expansion^6–8^ (GCE), in which orthogonal aaRS enzymes generate acyl-tRNAs that deliver non-canonical α-amino acids (ncAAs) to the ribosome. Once these acyl-tRNAs are accommodated within the A-site, the ribosome catalyzes incorporation of the ncAA at one or more sites within a protein. To date, hundreds of distinct ncAAs have been incorporated into proteins using orthogonal aaRS/tRNA pairs.^6,7^

Recent work has focused attention on expanding traditional genetic code expansion to include monomers that are not strictly α-amino acids. Examples of non-α-amino acid (nαAA) monomers that have been introduced into short peptides *in vitro* include expanded backbone amino acids (□^9–12^ or γ-^13,14^ amino acids, or derivatives of amino benzoic acid^15,16^) as well as monomers with distinct nucleophiles or functional groups in place of the α-amino group^17–23^. Incorporation of nαAA monomers into peptides *in vitro* benefits from the use of flexizymes: small structured RNAs that stoichiometrically acylate tRNA with diverse synthetic amino acid esters, including both ncAA as well as nαAA monomers^24^. Current flexizymes are not orthogonal and their activity in cells has not been demonstrated^24^. Thus, incorporation of nαAA monomers into proteins *in vivo* demands a suite of orthogonal synthetases that efficiently and selectively acylate tRNA in cells with the desired non-α-amino acid. The great current interest in sequence-defined hetero-polymers containing nαAA backbones^25,26^ demands multiple, mutually orthogonal synthetase enzymes that selectively activate these monomers and companion tRNAs that deliver them to wild type or modified ribosomes; success may also ultimately demand bespoke ribosomes and/or translation factors.

A number of methods are available to assess aaRS activity. A straightforward *in vivo* method is to simply assess the yield and purity of a protein whose mRNA contains a stop codon that is suppressed in the presence of a non-canonical amino acid and the corresponding orthogonal aaRS/tRNA pair. This method is most useful when the non-canonical amino acid is an α-amino acid, as there is greater confidence that the tRNA will be effectively delivered to the ribosome *in vivo* by the translation factor known as elongation factor thermo unstable (EF-Tu) and react efficiently once accommodated within the ribosomal A site. It is less useful for nαAA monomers, as differences in incorporation yield could result from differences in tRNA acylation efficiency, EF-Tu-promoted ribosome delivery^27^, peptide bond formation^28,29^, monomer uptake^30^, metabolism^17^, or other factors.

In our work on the identification of aaRS enzymes that accept non-α-amino acid substrates^31^, we sought a way to evaluate enzyme activity using purified proteins *in vitro*, bypassing the ribosome and other translation factors. We were most interested in a rapid, generalizable, and quantitative assay that directly evaluates the production of acyl-tRNA. To accommodate multiple aaRS enzymes and multiple nαAA substrates, the ideal assay should adapt easily to multiple enzymes, tRNAs, and non-α-amino acid monomers. Here we describe a convenient intact tRNA LC-MS assay that satisfies these requirements. tRNA is acylated *in vitro*, extracted, and analyzed via intact tRNA liquid-chromatography-mass spectrometry. This procedure provides the exact mass of the acyl-tRNA, quantifies reaction yields as low as 0.23%, and identifies acyl-tRNA products that cannot be distinguished from non-acyl-tRNAs using traditional gel electrophoresis.

### Development and overview of the protocol

In previous work, we made use of a degradative RNAse A-based assay to monitor formation of acylated tRNAs produced either by aaRS enzymes^31^ or flexizymes^16^. This assay detects the mass of the acylated 3’-terminal adenosine^32^ product that results from RNAse A-catalyzed degradation of the acyl-tRNA (RNAse A assay, Fig. 2a). Although useful for establishing *whether* a tRNA has been acylated, the RNAse A assay is less useful for establishing the precise yield of an aaRS-catalyzed reaction, especially when quantitative comparisons between monomers, tRNAs, and enzyme variants are required. This limitation arises because absolute quantification of tRNA acylation yield using the RNAse A assay would require the chemical synthesis of multiple acyl-adenosine standards, polar molecules with multiple chiral centers. Synthesis of these acyl-adenosine standards would undermine the generalizability of the assay. Because of the limitations of the RNAse A assay, we sought to develop an alternative, ratiometric mass spectrometry-based method to quantitatively evaluate the efficiency of aaRS-catalyzed tRNA acylation.

**Figure 2.**
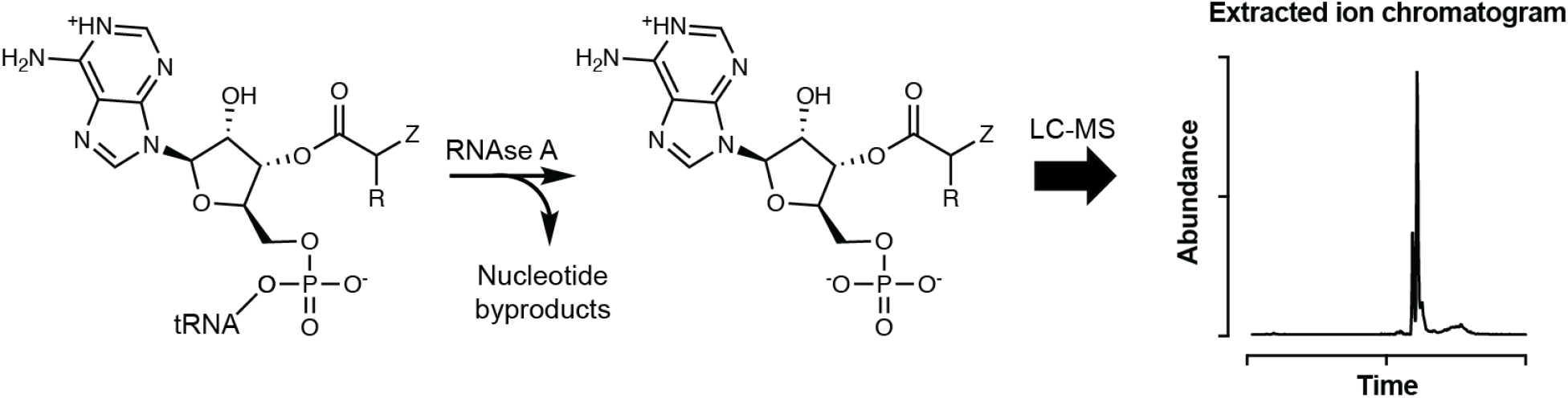
Scheme illustrating the mass-based detection of a unique acyl-adenosine after RNAse A-catalyzed tRNA degradation. Whether acylated or not, the 3’-adenosine is the only RNAse A product lacking a phosphate linkage. Two isobaric peaks are observed because acylation can occur on either the 2’- or 3’-hydroxyl group of the terminal adenosine and equilibration is rapid.

Intact tRNA molecules have been detected and analyzed using mass spectrometry for more than thirty years^33^. While there are some reports^34^ in which acyl-tRNA abundance is quantified using mass spectrometry, much of the tRNA MS literature uses mass spectrometry to identify post-transcriptional tRNA modifications as opposed to tRNA acylation^35^. That being said, there are several examples^36,37^ in which acyl-tRNAs are analyzed using matrix-assisted laser desorption/ionization mass spectrometry (MALDI-MS). In these examples, MALDI-MS was adapted for quantitative analysis using isotopically labeled substrates. We did not favor this method because the requirement for isotopically labeled substrates undermines generalizability.

We turned instead to LC-MS, which easily resolves acyl- and non-acyl-tRNA using ion-pairing reverse-phase chromatography and determines the exact masses of the species in each peak using high-resolution time-of-flight mass spectrometry. Because the non-acyl-tRNA peak in each total ion chromatogram (TIC) can contain tRNA species that cannot be enzymatically acylated (primarily due to substrate tRNAs that lack the 3’ terminal adenosine, a common byproduct of transcribing tRNAs *in vitro*^38^), simple integration of the acyl- and non-acyl-peaks in the A_260_ or total ion chromatograms may not accurately quantify the acylation yield. Therefore, we developed this method using the integration of extracted ion chromatograms to quantify the relative abundance of each tRNA species. Intact tRNA LC-MS offers exceptional analytical power, allowing for the exact mass detection of acyl-tRNA in addition to more unexpected species such as diacyl-tRNAs^39^ (in which both the 2’ and 3’ -OH of the 3’-terminal adenosine are acylated with the amino acid) as well as acylation of tRNAs that contain extra nucleotides, *vide infra*.

One problem we faced during the development of this intact tRNA acylation assay was the appearance of multiple cationic tRNA adducts that complicated analysis of mass spectra. These adducts were observed for both acyl- and non-acyl-tRNA and included adducts of sodium, potassium, and iron ions. Although there have been some reports^34,40^ that addition of ethylenediaminetetraacetic acid (EDTA) to an RNA sample prior to LC-MS analysis reduced cationic adduct formation, others^41^ have described conflicting results.

We found that the addition of EDTA was essential to eliminate tRNA-cation adducts. When no EDTA is used in the mobile phases, the base peak corresponding to *Methanomethylophilus alvus* pyrrolysyl-tRNA (*Ma*-tRNA^Pyl^, 22960 Da) in the deconvoluted mass spectrum can be an exceptionally small fraction (as little as 1%) of the observed masses (Fig. 3a). Even worse, the extent of adduct formation varied between injections and samples, severely complicating subsequent analysis. The majority of observed peaks correspond to deconvoluted masses of the base peak as an adduct with sodium, potassium, and/or iron. Upon addition of 27 µM EDTA free acid to the mobile phases of the LC-MS, the appearance of cation adducts was severely decreased in both the raw mass spectrum and the deconvoluted mass spectrum (Fig. 3c). However, at 27 µM EDTA, the raw mass spectrum contained many additional and unidentifiable peaks, presumably due to the overabundance of EDTA. When the concentration of EDTA in the mobile phases was decreased to 5 µM, the excess peaks in the raw mass spectrum were diminished, and there were few, if any, cation adducts observed in the deconvoluted mass spectrum (Fig. 3b).

**Figure 3.**
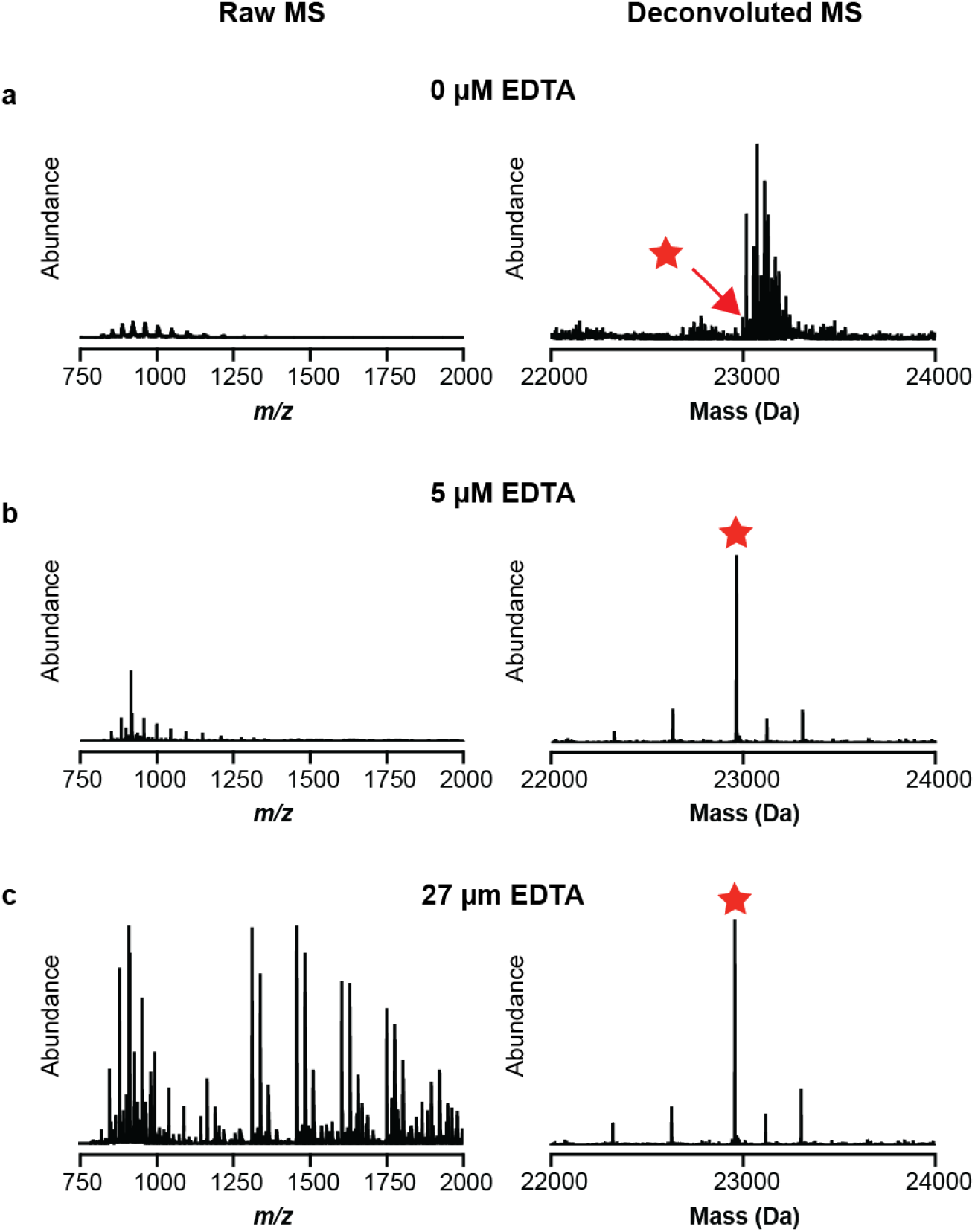
Mobile phases supplemented with EDTA eliminate confounding cation adducts of *Ma*-tRNA^Pyl^ during LC-MS analysis. Shown is the LC-MS analysis of pyrrolysyl-tRNA from *Methanomethylophilus alvus* (*Ma*-tRNA^Pyl^). The base peak (22960 Da) of the tRNA is identified by the red star. The concentration of EDTA in each panel is **a**, 0 µM, **b**, 5 µM, and **c**, 27 µM.

### Applications of the method

Acyl-tRNA can be obtained from acylation reactions performed either *in vitro* or *in vivo*. The *in vitro* acylation of tRNA is typically achieved using purified aaRS enzymes^42^, flexizyme-promoted reactions^24^, or, more historically, through chemoenzymatic means wherein a dinucleotide is first chemically acylated with the monomer of interest followed by ligation of the resulting acyl-dinucleotide to a tRNA lacking the two 3’-terminal nucleotides to form an acyl-tRNA^43^. Acyl-tRNA derived from any of these techniques can be analyzed using intact tRNA LC-MS as described in this protocol. While application of our protocol to tRNA generated *in vivo* is beyond the scope of this paper, methods to extract tRNA generated in cells have been reported^44^ and we envision it would be feasible with appropriate optimization. This protocol could also be modified to analyze other reactions producing covalently modified oligonucleotides^45–47^.

The intact tRNA LC-MS assay described here was employed in three recent studies. In one^31^, it quantified the relative activities of a suite of *M. alvus* PylRS variants for multiple canonical and non-canonical α-amino acids as well as non-α-amino acid substrates. Not only could the assay quantify the yields of acyl-tRNAs produced in aaRS-catalyzed reactions across multiple enzymes and multiple substrates, it also detected mono-acylation yields as low as 0.23% and revealed the relative propensity of each monomer/aaRS/tRNA triad to generate products that resulted from acylation twice, presumably at the 2’ and 3’-hydroxyl groups of the 3’-terminal ribose. In a second paper^48^, the intact tRNA LC-MS assay confirmed the identity of tRNA molecules acylated with various aminobenzoic acid derivatives before their use in intact ribosome cryo-EM studies. In this case, the tRNAs were unique because the 3’-terminal adenosine carried a 3’-NH_2_ substituent in place of the 3’-OH substituent. In more recent work, the intact tRNA LC-MS assay quantified the yield of flexizyme-promoted acylation reactions that were difficult to detect using gel electrophoresis. When using flexizymes to acylate tRNA, the yield is typically determined by acid-urea polyacrylamide gel electrophoresis (acid-urea PAGE)^24^. Here we show that intact tRNA LC-MS can detect activity for small, uncharged substrates that are not readily detected by acid-urea PAGE.

### Comparison with other methods

Many other methods have been reported to monitor aaRS activity and/or the production of acyl-tRNAs (Fig. 4, Table 1). Broadly, these methods fall into two categories: those that monitor the first chemical step in the reaction sequence (formation of the acyl-adenylate) and those that monitor the entire reaction (formation of the final acyl-tRNA reaction product).

**Figure 4.**
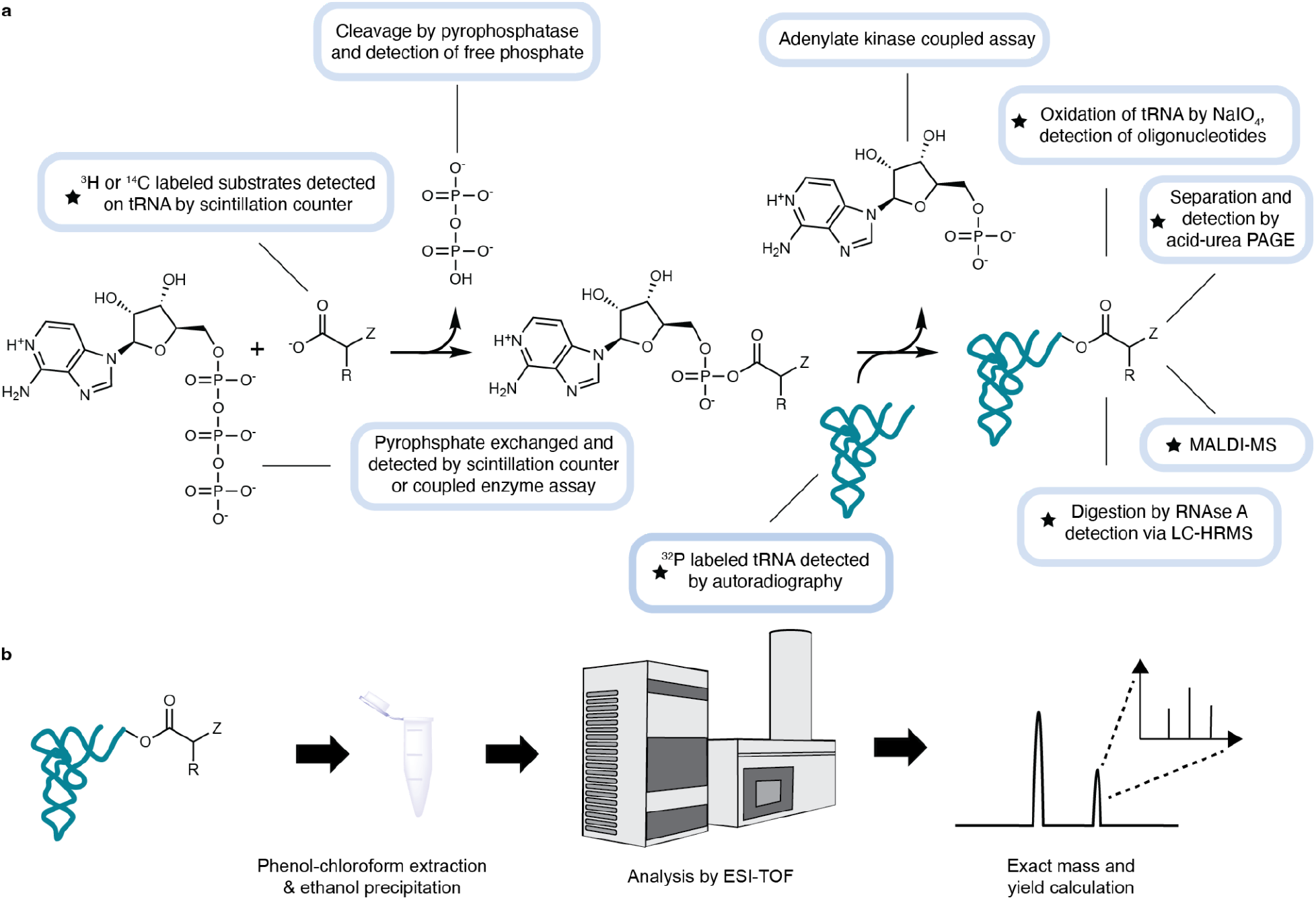
Schematic of different tRNA acylation assays. **a,** Overview of tRNA acylation by aaRSs and the different assays used to detect acylation. A star indicates the assay directly detects acyl-tRNA rather than a byproduct. **b,** schematic of intact tRNA LC-MS as described by this paper.

**Table 1.** Assays used to monitor aaRS-catalyzed reactions. a: ^32^P-labeled tRNA based techniques are generalizable with respect to the substrate and enzyme but not the tRNA. b: The RNAse A assay can be quantitative if acyl-adenosine standards are synthesized. c: MALDI-MS can be quantitative if isotopically labeled substrates are used.

| Assay | Step detected | Direct | Generalizable | Quantitative | Special requirements |
| --- | --- | --- | --- | --- | --- |
| Malachite green<sup>49</sup> | 1 | No | Yes | Yes |  |
| MesG<sup>50–52</sup> | 1 | No | Yes | Yes |  |
| <sup>32</sup>P pyrophosphate exchange<sup>53,54</sup> | 1 | No | Yes | Yes | Radioactive materials |
| AMP-PNP pyrophosphate exchange<sup>55</sup> | 1 | No | Yes | Yes |  |
| Adenylate kinase coupled assay<sup>56</sup> | 2 | No | Yes | Yes |  |
| Deacylase-modified AMP deaminase assay<sup>57–59</sup> | 2 | No | No | Yes |  |
| <sup>14</sup>C and <sup>3</sup>H labeled substrates<sup>60,61</sup> | 2 | Yes | No | Yes | Radioactive materials |
| <sup>32</sup>P labeled tRNA<sup>62,63</sup> | 2 | Yes | No<sup>a</sup> | Yes | Radioactive materials |
| Acid-urea PAGE<sup>64–67</sup> | 2 | Yes | No | Yes |  |
| IO<sub>4</sub> oxidation-oligo annealing<sup>68–70</sup> | 2 | Yes | Yes | Yes |  |
| RNase A assay<sup>16,32</sup> | 2 | Yes | Yes | No<sup>b</sup> | High resolution MS |
| MALDI-MS<sup>36,37</sup> | 2 | Yes | Yes | No<sup>c</sup> | High resolution MS |
| Intact tRNA LC-MS | 2 | Yes | Yes | Yes | High resolution MS |

Two widely used assays to indirectly detect acyl-adenylate formation use colorimetric monitoring of phosphate, derived from pyrophosphate generated during acyl-adenylate formation, upon reaction with either malachite green molybdate^49^ or 7-methylthioguanosine (MesG)^50–52^. These assays are employed widely because they are convenient, requiring only a spectrophotometer and relatively accessible reagents. The MesG assay, in particular, can be run continuously, another genuine advantage. Other indirect assays^53–55^ take advantage of the substrate-dependent exchange of pyrophosphate by the aaRS and quantify the ATP (or ATP analog) produced by this reaction. While these assays are convenient and widely used, they monitor only the first step in a two-step sequence, potentially missing structure-activity relationships that alter the rate or fidelity of the second step.

As mentioned previously, both RNAse A^16,32^ and MALDI-MS^36,37^ have been used to detect the complete aaRS-catalyzed reaction, but both techniques require the synthesis of internal standards to be quantitative, undermining their generalizability. Other strategies that monitor the complete aaRS-catalyzed reaction include a coupled enzymatic assay^56^ that relies on the release of AMP during the second step of the tRNA acylation reaction. This assay couples the activities of adenylate kinase, pyruvate kinase, and lactate dehydrogenase to the oxidation of NADH, which is detected spectrophotometrically. While free AMP should only be released following tRNA acylation by the acyl-adenylate intermediate, it is subject to false positive readings if the acyl-adenylate hydrolyzes spontaneously. Other coupled assays that detect the release of AMP through an enzyme cascade of AMP deaminase and IMP dehydrogenase to NADH/NAD^+^ redox chemistry have been reported^57–59^. These assays rely on enzymatic deacylation of the acyl-tRNA to regenerate non-acyl-tRNA and amplify the signal observed. However, the enzymatic deacylation requires an enzyme specific to that acyl-tRNA, and therefore these methods are not generalizable.

Early techniques^60,61^ to directly monitor the complete aaRS-catalyzed reaction relied on ^3^H and ^14^C radiolabeled substrates. However, radiolabeled amino acids are expensive, making it challenging, if not impossible, to achieve saturating concentrations. Moreover, it is unlikely that the radiolabeled version of more unique monomers are commercially available. This problem was resolved by the development of assays that employed ^32^P-labeled tRNA^62,63^, allowing for saturating concentrations of substrate. These ^32^P-labeled tRNA-based assays are rapid and can give precise quantitative data, but ^32^P is hazardous and requires special handling and authorizations.

Acid-urea PAGE exploits differences in electrophoretic mobility to separate^64–66^ acyl-tRNA from non-acyl-tRNA. This technique requires equipment and reagents that are common in laboratories; however, the degree to which electrophoretic mobility of acyl-tRNA differs from non-acyl-tRNA depends on monomer identity^64^ and can be very small, making the method prone to false negatives. In addition, even when electrophoretic mobility differences are sufficient, very long running times are required. In a modified acid-urea PAGE based method^67^, the acyl-tRNA is biotinylated at the α-amine, then bound to streptavidin before gel loading, improving the separation of acyl- and non-acyl-tRNA. However, biotinylation is not compatible with substrates that have modified α-substituents (nαAAs), which are of growing interest in the field of genetic code expansion.

Another method that detects intact tRNA relies on the fact that acylation of tRNA at the 2’- or 3’-hydroxyl group of the 3’-terminal adenosine protects the ribose from ribose oxidation by periodate^68–70^. Ribose oxidation inhibits ligation of the tRNA to an oligonucleotide. As a result, changes in the extent of ligation monitor the extent of tRNA acylation. In early work, non-oxidized tRNAs were ligated to a fluorescently labeled adapter oligonucleotide and fluorescence was measured via microarray^68^. Charged DM-tRNA-seq^69^ uses periodate oxidation followed by beta-elimination and next-generation sequencing to quantify tRNA acylation and can quantify tRNA charging of all cellular tRNAs. The most recent iteration, tREX (tRNA extension)^70^ uses a DNA polymerase to extend non-oxidized (and thus acylated) tRNA to form a fluorescently labeled RNA-DNA hybrid strand that can be detected via PAGE. These periodate-based techniques can report on acyl-tRNA derived from both *in vitro* and *in vivo* sources, but optimization of the periodate oxidation step is non-trivial, and unlike mass spectrometry-based methods, only report *if* a tRNA is acylated. They do not shed light on the identity of the monomer attached to tRNA, that is, whether it is the desired ncAA (or nαAA) or a natural α-amino acid.

While several of the assays described above are useful in different contexts, intact tRNA LC-MS has several distinct advantages. By detecting the exact mass of the acyl-tRNA, intact tRNA LC-MS offers the greatest analytical power of any method, comparable only to other MS-based methods^36,37^. Moreover, intact tRNA LC-MS is sensitive, rapid, quantitative, and generalizable with respect to the enzyme, substrate, and tRNA.

### Experimental design

This protocol describes the analysis and quantification of an *in vitro* tRNA acylation reaction via intact tRNA LC-MS. As such, it requires production and purification of both the tRNA and aaRS or the use of flexizyme. Some of the experiments described here were performed using PylRS from *Methanomethylophilus alvus* (*Ma*PylRS), which is easily expressed in *E. coli* and purified. PylRS enzymes from other species contain poorly soluble domains^71,72^, something that should be considered when choosing how to express and purify aaRSs. Similarly, the tRNA of interest must be purified, either following expression *in vivo* or transcription *in vitro*, and different tRNA sequences require different optimizations for production. *In vitro* transcription of some tRNA sequences can benefit from inserting a self-cleaving hammerhead ribozyme between a strong promoter and the tRNA gene to be expressed^73^. Specific variables of the *in vitro* tRNA acylation reaction including buffer composition and pH as well as the concentrations of the aaRS, tRNA, and substrate may need to be optimized. We also applied this method towards analyzing tRNA acylated via flexizyme, a technique that requires additional troubleshooting that has been discussed previously^24^. While this protocol describes the use of an LC-MS instrument from Waters Corporation, presumably the general protocol is applicable for instruments from other manufacturers. If using alternative instruments, it may be necessary to make modifications to the column, mobile phases, and method in order to obtain best results.

### Expertise needed to implement the protocol

RNA is susceptible to both base- and enzyme-catalyzed hydrolysis, and proper precaution must be taken to avoid degradation. RNAses are ubiquitous in the environment and can retain activity even after treatment in an autoclave^74^. Physical tools used for executing work with RNA such as this protocol should be treated with an RNAse inhibitor, such as RNase AWAY™ (Thermo Scientific) or RNaseZap™ (Invitrogen) prior to use. Reagents used for work with RNA should be separated from general use reagents to protect them from RNAse contamination. Familiarity with LC-MS would be beneficial for executing this protocol. If this protocol is not performed on an LC-MS dedicated to RNA work, an experienced LC-MS technician should evaluate how to best adapt the instrument for this protocol.

## Limitations

Intact tRNA LC-MS allows for rapid and quantitative analysis of tRNA acylation. While this protocol makes use of liquid chromatography and a reverse-phase column to resolve acyl-tRNA from non-acyl-tRNA, alternative solid supports and/or chromatography methods may be necessary for more complex product mixtures. Additionally, it is not certain how all aaRSs tolerate *in vitro* reactions, and optimization of the *in vitro* acylation conditions might be necessary for a given aaRS. Presumably this technique could be adapted to quantify tRNA acylation in tRNA that is expressed, charged, and purified from cells, but the current protocol does not describe that process. Given that tRNA from each acylation reaction must be extracted and precipitated and manual data analysis must be performed on each individual sample, this assay is not ideal for the most high-throughput applications. However, both of these limitations could be ameliorated through the use of an automated liquid handler and the development of software to automate the data analysis, respectively. For experiments that require more high-throughput analysis but less analytical power, alternative techniques such as charged DM-tRNA-seq might be better suited. This protocol requires a high resolution mass spectrometer, an instrument that is both expensive and requires sensitive upkeep.

## Materials

### Reagents

- HEPES: 4-(2-hydroxyethyl)-1-piperazineethanesulfonic acid (American Bio, cat. no. AB00892)
- Potassium hydroxide (Sigma Aldrich, cat. no. 221473)
- Magnesium Chloride, 1 M solution (American Bio, cat. no. AB09006)
- Adenosine 5’-triphosphate disodium salt hydrate (TCI, cat. no. A0157)
- Dithiothreitol (American Bio, cat. no. AB00490)
- *E. coli* inorganic pyrophosphatase (NEB, M0361)
- Phenol, saturated with citric acid, pH 4.5 (Millipore Sigma, cat. no. 6702-400ML)
- Chloroform (Avantor J.T. Baker, 925702)
- Sodium acetate 3.0M solution, pH 5.2 (American Bio, cat. no. AB13168)
- Ethylenediamine tetraacetic acid - EDTA (Fisher Scientific, cat. no. BP118)
- 1,1,1,3,3,3-Hexafluoro-2-propanol (Honeywell-Fluka, cat. no. 4206050ML)
- Triethylamine (Sigma Aldrich, cat. no. 90338-10X2ML)
- Methanol, Optima™ LC/MS Grade (Fisher Scientific, cat. no. A456-1)
- aaRS substrate - in the data shown below we used *N*^6^-(*tert*-butoxycarbonyl)-*L*-lysine (Millipore Sigma, cat. No. 359661-5G)
- Ethanol (Millipore Sigma, cat. No. E7023-500ML)
- Bicine: *N,N*-Bis(2-hydroxyethyl)glycine (Millipore Sigma, cat. No. B3876-25G)
- γ-ketoester flexizyme monomer: 3,5-dinitrobenzyl 4-oxopentanoate (synthesized in accordance with Lee et al.^75^

### Biological materials

- Purified aaRS (procedure described in our previous study^31^)
- Purified tRNA (procedure described in our previous study^31^)
- dFx (Flexizyme - Integrated DNA Technologies. Sequence: GGAUCGAAAGAUUUCCGCAUCCCCGAAAGGGUACAUGGCGUUAGGU)
- MH (Microhelix tRNA mimic - Integrated DNA Technologies. Sequence: GGCUCUGUUCGCAGAGCCGCCA)

### Equipment

- Analytical balance (Mettler Toledo, cat. no. XP26)
- Micropipettes: p2, p10, p200, p1000 (Rainin, cat. nos. 17014413, 17014409, 17014411, 17014407)
- Pipette tips (USA Scientific cat. nos. 1111-3810, 1110-1810, 1111-2830)
- 1.5 mL microcentrifuge tubes (Fisher Scientific, cat. no. 05-408-129)
- Refrigerated microcentrifuge (Eppendorf, cat. no. 5425 R)
- -80 °C freezer (Thermo Scientific, cat. no. TSX60086A)
- Milli-Q® Advantage A10 Water Purification System (Millipore Sigma, cat. no. Z00Q0V0WW)
- ACQUITY UPLC I-Class PLUS (Waters, cat. no. 186015082)
- ACQUITY UPLC BEH C18 Column, 130 Å, 1.7 µm, 2.1 mm X 50 mm (Waters, cat. no. 186002350)
- Xevo G2-XS Tof (Waters, cat. no. 186010532)
- Autosampler vials, 350 µL fused glass insert (Thermo Scientific, cat. no. C4011-LV1W)
- Autosampler snap-it caps (Thermo Scientific, cat. no. C4011-50)
- PYREX Corning 500mL Round Media Storage Bottles (PYREX Corning, cat. no. 1395-500)
- Ultrasonic Cleaner 2L (Vevor, cat. no. B01HGN40SQ)

### Reagent Setup

#### Preparation of mobile phases

  - CRITICAL: Use brand new, perfectly clean glassware for preparation of mobile phases and dedicate this glassware for that purpose. The glassware should never be exposed to detergents to prevent contamination on the column.
  - CRITICAL: The UPLC solvent lines should also be used exclusively for the mobile phases and not other solvents.
  - CRITICAL: In this section, “water” refers to fresh ultrapure water with 18.2 MΩ resistivity (ie. water from a MilliQ Water Purification System).

- Clean bottles using the following procedure before use: rinse bottles with water three times, then fill bottle completely with water, sonicate the bottle for 5 minutes at room temperature, discard the water. The bottle is now ready to be used.
  - CAUTION: preparation of mobile phases should be performed in a chemical fume hood while wearing proper personal protective equipment (nitrile gloves, laboratory coat, and safety glasses).
- Prepare the 500 µM EDTA solution. Tare a 500 mL borosilicate glass bottle. Add 73 mg of EDTA free acid and 500 g of water. Place in a sonicator/heat bath and sonicate at 50 °C for 4 hours. Note that EDTA requires a long time to dissolve.
  - CRITICAL: EDTA is added to the mobile phases to scavenge metal cations. EDTA is most frequently sold and stocked as a sodium-containing buffered solution. Because the purpose of the EDTA is to remove cations, EDTA free acid should be used, not the sodium-containing buffered solution.
- Prepare Mobile Phase A. Tare a 500 mL borosilicate glass bottle. Add 490.2 g water, 5 mL 500 µM EDTA solution, 557.6 µL triethylamine (TEA), and 4.2 mL of hexafluoroisopropanol (HFIP). Mix by inversion several times and sonicate the bottle for 5 minutes.
  - CRITICAL: To ensure optimum quality mobile phases, a fresh ampoule of triethylamine should be used.
  - CAUTION: HFIP is corrosive and hazardous. Work with HFIP should be performed in a chemical fume hood while wearing personal protective equipment (nitrile gloves, laboratory coat and safety glasses).
- Prepare Mobile Phase B. Tare a 500 mL borosilicate glass bottle. Add 242.6 g water, 250 mL methanol, 5 mL 500 µM EDTA solution, 278.8 µL TEA, and 2.1 mL HFIP. Mix by inversion several times and sonicate the bottle for 5 minutes.
  - CRITICAL: Mobile phases should be prepared fresh frequently; we recommend replacing them every two weeks or at the most four.

## Procedure

### tRNA acylation reaction - 2-25 hours

1. For a 25 µL reaction, add the following components to a 1.5 mL microcentrifuge tube:
  a. 100 mM Hepes-KOH (pH 7.5)
  b. 10 mM MgCl_2_
  c. 10 mM ATP
  d. 4 mM DTT
  e. 0.1 U *E. coli* inorganic pyrophosphatase
  f. 0-10 mM substrate
  g. 25 µM tRNA
  h. 2.5 µM aminoacyl-tRNA synthetase (more can be used if the aaRS/substrate pair has low activity)
  i. Water to 25 µL
2. Mix components well, spin down in a microcentrifuge to ensure all materials are at the bottom of the tube, and incubate at 37 °C for 1-24 hours (2-6 hours is recommended).

### Phenol-chloroform extraction and ethanol precipitation of acylation reaction - 1.5-2 hours (depending on number of samples)

3. Add 20 µL sodium acetate buffer (3 M, pH 5.2) and 155 µL water to each sample to a final volume of 200 µL.
4. Add 100 µL phenol and 100 µL chloroform. Mix vigorously for 30 seconds. Spin in microcentrifuge at 21,300 x g for 30 seconds at 4 °C to separate the phases. Remove the organic phase from the bottom using a 200 µL micropipette.
  a. CRITICAL: To minimize the hydrolysis of the acyl-tRNA, this extraction as well as other sample manipulation should be performed on ice.
5. Repeat extraction (Step 4) twice more with just 200 µL chloroform and no phenol.
6. To each 200 µL aqueous solution of acyl-tRNA following phenol chloroform extraction, add 500 µL of 100% ethanol and shake well to mix.
7. Place microcentrifuge tubes in a -80 °C freezer for 30 minutes to precipitate tRNA.
8. Transfer microcentrifuge tubes to a microcentrifuge and spin at 21,300 x g for 30 minutes at 4 °C. There should be a very small (∼ < 1 mm) pellet of tRNA at the bottom of the microcentrifuge tube. Remove supernatant and place tubes in a chemical fume hood for 10 minutes with the caps open to dry any remaining ethanol.
  a. CRITICAL: Be careful not to disturb the pellet when removing the supernatant.
  b. PAUSE POINT: tRNA pellets can be stored at -80 °C overnight or longer.

### Sample preparation - 0.5 hours

9. Dissolve the pellet in 2.25 µL water. Add 19 µL water to an autosampler vial with insert. Transfer 1 µL of tRNA solution to the autosampler vial and mix. Assuming >99% recovery of tRNA during the workup, the tRNA concentration should be approximately 13 pmol/µL.
  a. Because small molecules from the acylation reaction can remain with the tRNA following the extraction and precipitation that absorb in the UV range, measuring the concentration of the tRNA by Nanodrop is not accurate at this point.
  b. If the instrument used for analysis has a blunt needle, be sure to use pre-slit caps or use a razor blade to cut a slit into the cap of the autosampler vial.

### Acylation, Ethanol Precipitation, and Sample Preparation for Flexizyme/Microhelix reactions

a. 3,5-dinitrobenzyl 4-oxopentanoate was acylated onto microhelix RNA following standard protocols for flexyzme acylation^24^. 1 μL of 250 μM Flexizyme (dFx or eFx) was added to 1 μL of 500 mM HEPES-KOH (pH 7.5) or 500 mM Bicine-KOH (pH 9.0) and 1 μL of 250 μM microhelix tRNA. The sample was incubated at 95°C for 2 min and allowed to reach room temperature in 5 min. 2 μL of 3 M magnesium chloride was then added, followed by 1 μL of a DMSO solution of 3,5-dinitrobenzyl 4-oxopentanoate (50 mM). Water was then added to 10 µL. The reaction was incubated at 4°C for 24 h.
b. Following Flexizyme acylation, samples were ethanol precipitated (70% EtOH, 0.1M sodium acetate, pH 5.2) and washed twice with 70% ethanol at 4°C to remove residual salt that could damage the MS or interfere with column chromatography. Samples were resuspended into 10 µL of water. 1 µL of this was injected (∼25 pmol) onto the LC-MS.

### LC-MS injection - 15 minutes per sample

10. Use the following LC method for separation of acyl-tRNA from non-acyl-tRNA
  a. Flow rate: 0.3 mL/min
  b. Initial condition: 22% mobile phase B
  c. Linear gradient over 10 minutes to 40% mobile phase B
  d. Linear gradient over 1 minute to 60% mobile phase B
  e. Linear gradient over 0.1 minutes to 22% mobile phase B
  f. Hold over 2.9 minutes at 22% mobile phase B
  g. Column temperature: 60 °C
11. Use the following MS method for detection of oligonucleotides
  a. Ion mode: negative
  b. Capillary voltage: 2000 V
  c. Sampling cone: 40
  d. Source offset: 40
  e. Source temperature: 140 °C
  f. Desolvation temperature: 20 °C
  g. Cone gas flow: 10 L/h
  h. Desolvation gas flow: 800 L/h
  i. Spectrum reads: 1/s
  j. Mass range: 500 - 2000 *m/z*
12. Inject tRNA sample. We recommend injecting 1 µL or approximately 13 pmol, though we observe usable results with injections between 5 and 20 pmol, or as much as 100 pmol for exceptionally low yielding substrates.

### LC-MS data analysis - 5 minutes per sample

Deconvolution of intact tRNA mass

a. The following protocol applies directly to the MassLynx software from Waters Corporation, but should be adaptable to other mass analysis software.

13. Open the total ion and UV chromatograms for your sample.
14. Identify the peaks that you want to analyze.
  a. Most, though not necessarily all, tRNA species elute between approximately 5 and 8 minutes.
  b. There is usually a large peak that elutes at ∼0.5 minutes that appears when any sample is injected. This is likely the small molecules present in the sample. To avoid wear on the detector with small molecules and salts, it is possible to divert the LC flow to to waste for the first 0.8 minutes of a run.
15. In the TIC (Fig. 6b), combine the mass spectra data for the peak you wish to analyze.
16. Zoom into the raw mass spectrum (Fig. 6c) and identify the peaks that correspond to your oligonucleotide. We refer to this range in the raw mass spectrum as the deconvolution range (Fig. 6d).
  a. Most tRNA charge-state envelopes will appear in the 800 - 1300 *m/z* range, though we frequently see them all the way up to 2000, and occasionally slightly below 800. tRNA charge-state envelopes contain multiple peaks corresponding to the isotopic distribution of molecules within that envelope. Identify the most abundant charge-state envelope corresponding to the tRNA species of interest. We call this the “major ion.”
17. Zoom in on the major ion (Fig. 6e) so you can identify the *m*/*z* of the peak that sits in the center of the charge-state envelope.
18. Measure the width of the major ion charge-state envelope and record this value as the width at half height.
19. Zoom into an *m/z* window where all of the tRNA charge-state envelopes are in frame, but you are excluding the *m/z* range that does not contain tRNA charge-state envelopes, the raw MS deconvolution range (Fig. 6d).
20. Deconvolute the mass of your tRNA species (Fig. 6f) using the MaxEnt1 plugin (or comparable deconvolution software). We recommend the following parameters:
  a. Output Mass Ranges: we recommend a range a few thousand below and above your expected mass, i.e. for a tRNA with expected mass of 24,000, use 21,000 - 27,000.
  b. Output Mass Resolution: we recommend 0.10 Da/channel. For faster calculations, use 0.20 Da/channel.
  c. Damage model: Uniform Gaussian. Enter the width at half height recorded earlier.
  d. Minimum intensity ratios: Left 33, Right 33.
  e. Completion options: Iterate to convergence.
21. Record the deconvoluted mass of your tRNA species.
  a. To calculate the expected mass of each tRNA species, we used an online RNA molecular weight calculator^77^ then calculated the additional molecular weight of the monomer of interest using ChemDraw 21.0.0.
22. Repeat this process for all tRNA species of interest including your acyl-tRNA peak and non-acyl-tRNA peak.

**Figure 5.**
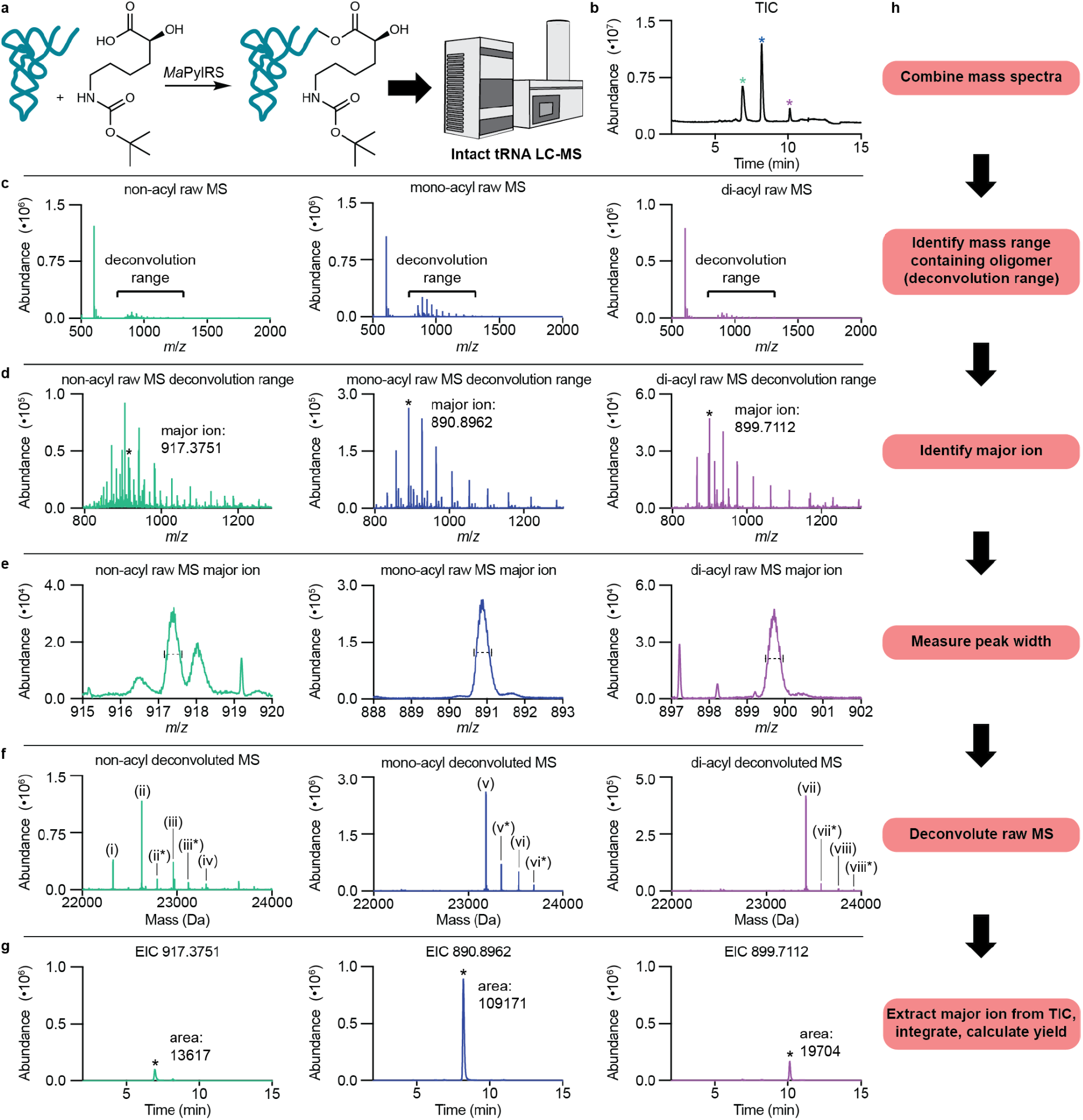
LC-MS analysis of a tRNA acylation reaction. **a**, Scheme of acylation reaction and analysis. **b**, Total ion chromatogram of the injection. Each species identified in the TIC is noted by a colored asterisk: non-acyl-tRNA (green), mono-acyl-tRNA (blue), and di-acyl-tRNA (purple). The color coding used in the TIC is also used in panels c-g. **c**, Raw mass spectra for each tRNA species. The mass range containing the oligonucleotide ladder of the RNA species of interest (deconvolution range) is used for the final deconvolution of the intact tRNA. This is usually approximately 800-1300 *m/z*. **d**, Raw mass spectra for each tRNA species zoomed into the oligonucleotide ladder. Here the most abundant charge state ion (major ion) corresponding to the tRNA species of interest can be identified. **e**, Raw mass spectra for each tRNA species zoomed into the major ion peak. For each major ion peak the width at half maximum can be measured for input in the deconvolution software. **f**, Deconvoluted mass spectra for each tRNA species. Each unique tRNA species identified is denoted by a Roman numeral in parentheses and described in Table 2. Multiple peaks are present in the deconvoluted mass spectrum of the non-acyl-tRNA, which is why simple integration of the TIC may not accurately quantify the acylation yield. **g**, Extracted ion chromatograms for the major ion of different tRNA species used for quantification. **h**, Flow chart of the data workup process.

**Figure 6.**
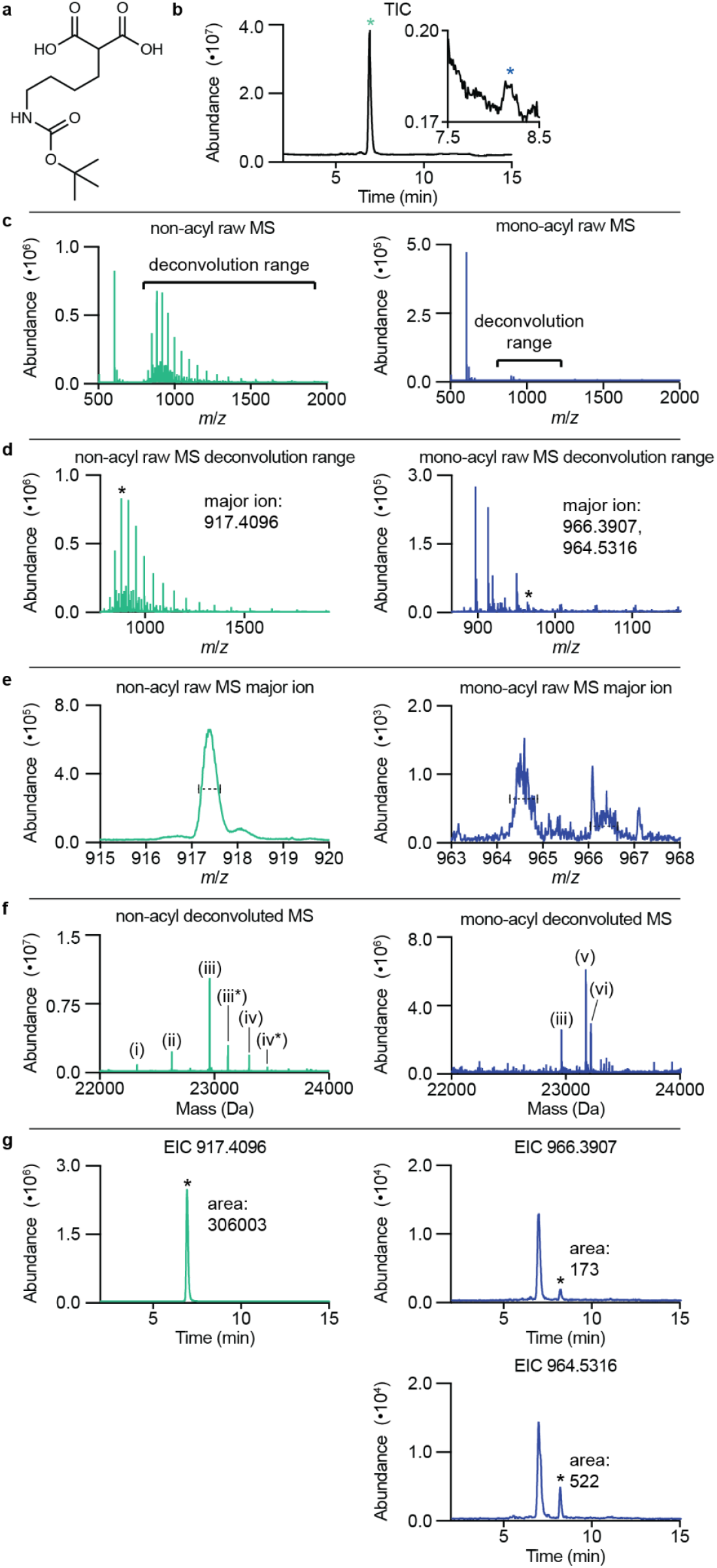
LC-MS analysis of a tRNA acylation reaction with an exceptionally low yield. **a**, Structure of the substrate for this experiment, BocK-malonate **b**, Total ion chromatogram of the injection. Each species identified in the TIC is noted by a colored asterisk: non-acyl-tRNA (green) & mono-acyl-tRNA (blue). The color coding used in the TIC is also used in panels c-g. **c**, Raw mass spectra for each tRNA species. The mass range containing the oligonucleotide ladder of the RNA species of interest (deconvolution range) is used for the final deconvolution of the intact tRNA. **d**, Raw mass spectra for each tRNA species zoomed into the oligonucleotide ladder. Here the most abundant charge state ion (major ion) corresponding to the tRNA species of interest can be identified. **e**, Raw mass spectra for each tRNA species zoomed into the major ion peak. For each major ion peak the width at half maximum can be measured for input in the deconvolution software. **f**, Deconvoluted mass spectra for each tRNA species. Each different tRNA species identified is denoted by a Roman numeral in parentheses and described in Table 3. Multiple peaks are present in the deconvoluted mass spectrum of the non-acyl-tRNA, which is why simple integration of the TIC may not accurately quantify the acylation yield. **g**, Extracted ion chromatograms for the major ion of different tRNA species used for quantification.

### Determination of ratios of tRNA species in sample

Because the non-acyl-tRNA peak in each total ion chromatogram (TIC) contains tRNA species that cannot be enzymatically acylated (primarily tRNAs that lack the 3’ terminal adenosine^38^), simple integration of the acylated and non-acylated peaks in the A_260_ or total ion chromatograms may not accurately quantify the acylation yield. Use the following procedure to accurately quantify the ratio of acyl-versus non-acyl-tRNA.

23. Extract the major ion corresponding to each tRNA species of interest (Fig. 6g). For the mass chromatogram window, we use ± 0.3000 Da, as this is the approximate width at half height of most samples.
  a. CRITICAL: make sure that the peak in the extracted ion chromatogram (EIC) aligns with the peak from the TIC that contains your tRNA species.
24. Integrate the EIC.
25. Visually inspect the boundaries of the peak identified by the integration. Occasionally the automatic integration will need to be manually adjusted.
26. Record the area of the peak as determined by integration.
27. Repeat this process for all tRNA species of interest.
28. Calculate the yield of the tRNA acylation reaction using the ratios of the peak areas for the different tRNA species in the sample:

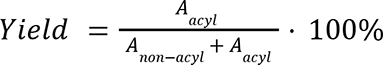

### Troubleshooting

| Step | Problem | Possible Cause | Solution |
| --- | --- | --- | --- |
| 14 | Multiple peaks present in chromatogram before main tRNA peak | Degradation of tRNA | Make sure to keep tRNA on ice except during enzymatic reactions, use RNase-free water, clean surfaces with RNase inhibitor solution |
| 14 | No peaks at all | Sample prep | tRNA not concentrated enough, ethanol precipitation failed, sample too salty |
| 14 | No acylated peak present in chromatogram | Low acylation reaction yield | Look for small peak on the shoulder of non-acyl-tRNA peak |
| 16 | Mass spectrum shows evidence of cation adducts | Cation concentration in system too high | Increase EDTA concentration in mobile phases |
| 16 | Mass spectrum contains many small molecule peaks | EDTA concentration is too high | Decrease EDTA concentration in mobile phases |

### Timing

1. Acylation reaction 2-25 hours (depending on aaRS & substrate activity)
2. Phenol-chloroform extraction and ethanol precipitation of samples 1.5-2 hours depending on number of samples
3. LC-MS injection - 15 minutes per sample
4. LC-MS data analysis - 5 minutes per sample

### Anticipated Results

In our previous study^31^, we used the intact tRNA LC-MS assay described herein to evaluate the ability of several *Methanomethylophilus alvus* PylRS variants to charge their cognate tRNA with a number of nαAAs. Here we describe one representative data set from that study (Fig. 6). In this experiment, wild type *M. alvus* PylRS was used to charge *M. alvus* tRNA^Pyl^ (*Ma-*tRNA^Pyl^) with *(S)*-6-((tert-butoxycarbonyl)amino)-2-hydroxyhexanoic acid (BocK-OH) (Fig. 6a). In this data set, the acyl-tRNA yield is shown to be 90.4%. The deconvoluted masses of different tRNA species are shown in Table 2.

**Table 2.** Deconvoluted masses of different tRNA species. The tRNA species identified in Fig. 6f are described here. Transcript length N refers to the correct tRNA transcript, while N-2 and N-1 refer to transcripts that lack two and one of the 3’ nucleotides, respectively, and N+G refers to a transcript whose mass corresponds to the correct transcript with the additional mass of one guanosine residue. 5’-phosphorylation refers to whether the 5’-terminal nucleotide contains a monophosphate or triphosphate. Acylation refers to whether the 3’-terminal adenosine is acylated with the monomer of interest zero, one, or two times.

| Label | Transcript length | 5'-phosphorylation | Acylation | Expected mass | Observed mass |
| --- | --- | --- | --- | --- | --- |
| (i) | N-2 | monophosphate | non- | 22326 | 22325.5 |
| (ii) | N-1 | monophosphate | non- | 22631 | 22630.7 |
| (ii*) | N-1 | triphosphate | non- | 22791 | 22790.8 |
| (iii) | N | monophosphate | non- | 22960 | 22959.9 |
| (iii*) | N | triphosphate | non- | 23120 | 23119.8 |
| (iv) | N+G | monophosphate | non- | 23305 | 23304.8 |
| (v) | N | monophosphate | mono- | 23189.4 | 23189.1 |
| (v*) | N | triphosphate | mono- | 23349.4 | 23349.3 |
| (vi) | N+G | monophosphate | mono- | 23534.4 | 23534.4 |
| (vi*) | N+G | triphosphate | mono- | 23694.4 | 23694.4 |
| (vii) | N | monophosphate | di- | 23418.8 | 23418.5 |
| (vii*) | N | triphosphate | di- | 23578.8 | 23578.6 |
| (viii) | N+G | monophosphate | di- | 23763.8 | 23766.8 |
| (viii*) | N+G | triphosphate | di- | 23923.8 | 23924.4 |

**Table 3.** Deconvoluted masses of different tRNA species. The tRNA species identified in Fig. 6f are described here. Transcript length N refers to the correct tRNA transcript, while N-2 and N-1 refer to transcripts that lack two and one of the 3’ nucleotides, respectively, and N+G refers to a transcript whose mass corresponds to the correct transcript with the additional mass of one guanosine residue. 5’-phosphorylation refers to whether the 5’-terminal nucleotide contains a monophosphate or triphosphate. Acylation refers to whether the 3’-terminal adenosine is acylated with the monomer of interest zero, one, or two times. (v) & (vi) are both acylated with BocK-malonate, but (v) is the decarboxylation product of the malonyl-tRNA.

| Label | Transcript length | 5'-phosphorylation | Acylation | Expected mass | Observed mass |
| --- | --- | --- | --- | --- | --- |
| (i) | N-2 | monophosphate | non- | 22326 | 22325.6 |
| (ii) | N-1 | monophosphate | non- | 22631 | 22630.7 |
| (iii) | N | monophosphate | non- | 22960 | 22960.1 |
| (iii*) | N | triphosphate | non- | 23120 | 23120.3 |
| (iv) | N+G | monophosphate | non- | 23305 | 23305.1 |
| (iv*) | N+G | triphosphate | non- | 23465 | 23465.3 |
| (v) | N | monophosphate | mono-decarboxylation | 23217.4 | 23217.2 |
| (vi) | N | monophosphate | mono-malonyl | 23173.4 | 23172.8 |

This data set highlights the high analytical power of intact tRNA LC-MS. In the deconvoluted mass spectrum for non-acyl-tRNA (Fig. 6f, left) multiple peaks are present. The expected transcription product, *Ma-*tRNA^Pyl^, is present both as the 5’ monophosphate and triphosphate forms—(iii) & (iii*). Some of the other peaks observed are early termination transcription products that lack either the 3’-terminal adenosine—(ii), & (ii*)—or both the 3’-terminal adenosine and penultimate cytosine residue—(i). Another peak—(iv)—corresponds to a transcription product that contains an additional guanosine, dubbed N+G. Because *Ma-*tRNA^Pyl^ initiates transcription with five consecutive 5’ guanosines, we presume that the additional guanosine is inserted at the 5’ end of the tRNA.

In the deconvoluted mass spectrum for the mono-acyl-tRNA (Fig. 6f, center) four species are identified corresponding to *Ma-*tRNA^Pyl^ acylated with one BocK-OH monomer. The correct transcription product—(v) & (v*), the 5’ mono and triphosphate versions of *Ma-*tRNA^Pyl^—were acylated with one equivalent of BocK-OH. Surprisingly, we found that the N+G tRNA species—(vi) & (vi*) was also acylated with one equivalent of BocK-OH. To the best of our knowledge, it has not been previously reported that tRNAs with additional nucleotides can be active in enzymatic acylation reactions, perhaps something that has gone previously unobserved due to the use of alternative analytical techniques.

In the deconvoluted mass spectrum for the di-acyl-tRNA (Fig. 6f, right), four species of tRNA are observed. Like in the mono-acyl-tRNA deconvoluted mass spectrum, both the correct transcription product and N+G product (monophosphate and triphosphate versions of each) are acylated with BocK-OH, in this case two equivalents. Di-acylation of tRNAs has been reported before^39^ and because it has been shown that di-acyl-tRNAs are active in translation^78^, we counted di-acyl-tRNAs toward the overall acylation yield calculations.

This data set also demonstrates why simple integration of the A_260_ or TIC may not accurately quantify the acylation yield. In the non-acyl-tRNA deconvoluted mass spectrum, some of the most abundant peaks—(i), (ii), & (ii*)—were of the early termination products that lack the 3’ terminal adenosine required for acylation activity. Because these tRNAs are not active acylation substrates for PylRS, they should not be counted when calculating the yield of the acylation reaction, but because they were not resolved from the active tRNA species, we must extract the active tRNA species in order to accurately calculate the acylation yield. Using the integrated areas of the extracted ion chromatograms (Fig. 6g), we can calculate that the total acylation yield:

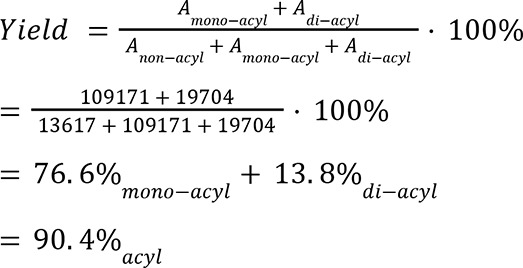

In our previous study^31^ we also demonstrated that this technique is capable of detecting acyl-tRNAs from reactions of enzyme/substrate pairs with exceptionally low yields. In this experiment, wild type *M. alvus* PylRS was used to charge 2-(4-((tert-butoxycarbonyl)amino)butyl)malonic acid (BocK-malonate, Fig. 6a) onto *Ma-*tRNA^Pyl^ and the yield was determined to be 0.23%. Upon first glance at the total ion chromatogram (Fig. 6b), it appears that there is only one peak corresponding to the non-acyl-tRNA and no mono-acyl-tRNA peak. However, when we zoomed into the shoulder of the non-acyl-tRNA peak, we were able to identify an additional peak that deconvoluted to the expected masses of the tRNA acylated with BocK-malonate as well as the decarboxylation product of this tRNA (Fig. 6f). We previously^31^ determined that the decarboxylation of this acyl-tRNA product occurs during the reaction workup or intact tRNA LC-MS analysis and therefore to calculate the overall acylation yield with BocK-malonate, we extracted the major ions for both the malonyl product as well as the decarboxylation product (Fig. 6g). The total yield of this reaction is calculated below:

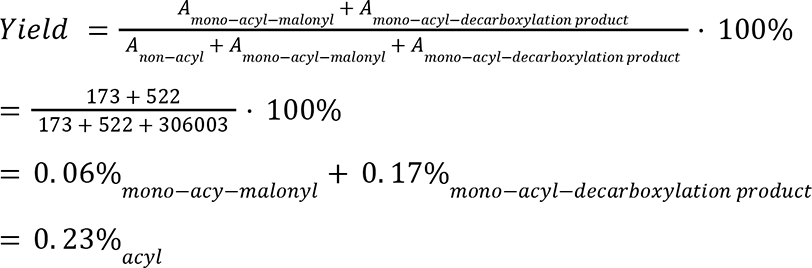

Intact tRNA LC-MS can also be used to quantify the acylation yields of flexizyme-mediate acylations. Traditional methods to quantify acylation efficiencies of flexizyme acylations rely on acid-urea PAGE to separate a non-acyl and mono-acyl microhelix RNA, a tRNA acceptor stem analogue. In cases in which the monomer is small or not positively charged, separation by gel electrophoresis can be challenging. Recent attempts to determine the flexizyme acylation of a small 4-oxopentanoate ester exemplify this problem: no separation was distinguishable and an acylation yield was indeterminable when using acid-urea PAGE^75^. We carried out the same flexizyme acylation of 4-oxopentanoate ester onto a microhelix and showed that our intact tRNA LC-MS methods can be used to resolve, confirm, and quantify the acylation of this small, uncharged monomer (Fig. 7).

**Figure 7.**
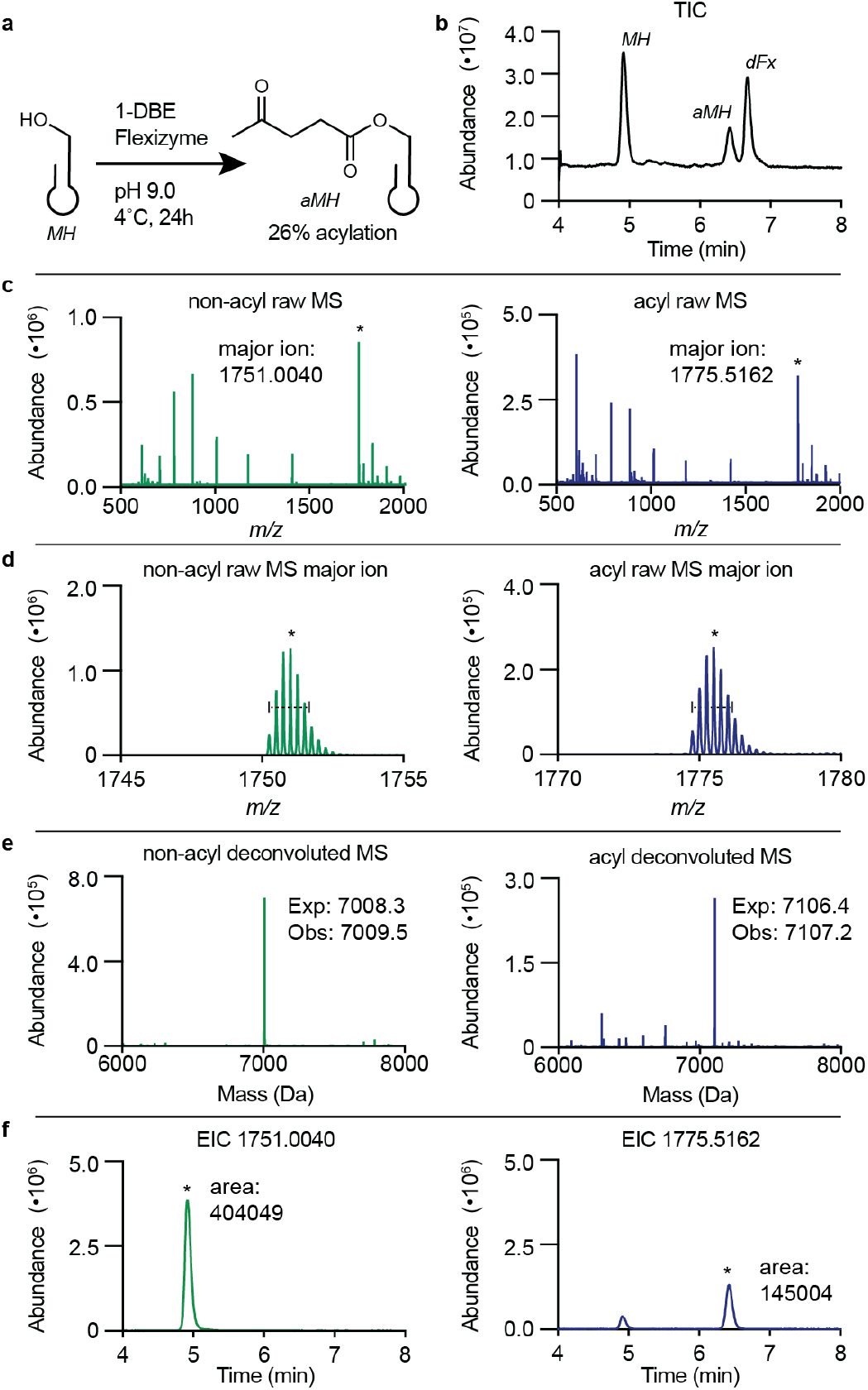
Intact tRNA LC-MS for the acylation of small, uncharged flexizyme monomers with microhelix. **a,** Representation of the acylation of microhelix (MH) with 3,5-dinitrobenzyl 4-oxopentanoate (1-DBE) to make the acylated microhelix (aMH). **b,** Total ion chromatogram (TIC) of the flexizyme reaction displaying the MH, aMH, and flexizyme (dFx) For the non-acyl-MH (left, green) and acyl-MH (right, blue), the raw mass spectra **c**, zoomed in views of the -4 charge state envelopes in the mass spectra **d**, Deconvoluted mass spectra for each RNA species **e**, and the extracted ion chromatograms (EIC) of the major ions **f** are shown.

## Author Contributions

R.F., I.K., and A.S. conceived of this project and prepared this manuscript. Initial optimization of this protocol and its application towards the analysis of enzymatic tRNA acylation reactions using aminoacyl-tRNA synthetases was performed by R.F.. Adaptation of this protocol towards analyzing tRNA acylation reactions using flexizyme was performed by I.K..

## Acknowledgements

We would like to thank Cameron V. Swenson for his helpful contributions and commentary towards this manuscript. This work was supported by the NSF Center for Genetically Encoded Materials, CHE-2002182.

